# Electrical coupling within thalamocortical networks cumulatively reduces cortical correlation to sensory inputs

**DOI:** 10.1101/2025.08.21.671565

**Authors:** Austin J. Mendoza, Julie S. Haas

## Abstract

Thalamocortical (TC) cells relay sensory information to the cortex, as well as driving their own feedback inhibition through collateral excitation of the thalamic reticular nucleus (TRN). The GABAergic cells of the TRN are extensively coupled through electrical synapses. While electrical synapses are most often noted for their roles in synchronizing rhythmic forms of neuronal activity, they are also positioned to modulate responses to transient information flow across and throughout the brain, although this effect is seldom explored. Here we sought to understand how electrical synapses embedded within a network of TRN neurons regulate the processing of ongoing sensory inputs during relay from thalamus to cortex. We used Hodgkin-Huxley point models to construct a network of a 9 TC and 9 TRN cells, with one cortical output neuron summing the TC activity. Each pair of TC and TRN cells was reciprocally coupled by chemical synapses. TRN cells were each electrically coupled to two neighboring cells, forming a ring topology. TC cells received synaptic inputs in sequence, with intervals between inputs varying from 10 to 50 ms across simulations. This architecture and sequence of inputs allowed us to assess the functional radius of an electrical synapse by comparing the cumulative effects of each additional TRN electrical synapse on the responses of the TRN and TC cells and the cortical output. Effects of electrical synapses on TRN cell activity were strongest for smaller intervals between inputs, and cumulative with additional synapses. In contrast, effects in TC neurons were strongest for larger intervals between inputs and also increased with coupling strength. Coupling within TRN modulated cortical integration of TC inputs by unexpectedly increasing response rates, duration and reducing spike correlation to the input sequence that was presented to the TC layer. Thus, embedded TRN electrical synapses exert powerful influence on thalamocortical relay, in a cumulative manner. These results highlight the multi-synaptic influences of electrically coupled cells and reinforce that they should be included in more complex and realistic networks of the brain.

## Introduction

Sensory information from the environment is communicated to the cortex by relay neurons of the thalamic nuclei. Rather than simple relay, however, it is generally agreed that neural information is transformed by thalamic circuitry. Ascending sensory information from all sensory systems, except for olfaction, targets cells of thalamic nuclei during transduction to higher brain regions (Sherman and Guillery, 2006; Jones, 2007). Thalamocortical (TC) cells within the primary sensory thalamic nuclei –the lateral geniculate nucleus for visual relay (Bickford, 2019), the ventral postmedial nucleus for touch (O’Reilly et al., 2021), the medial geniculate body for auditory inputs (Bartlett, 2013) -- receive strong “driver” inputs from sensory afferent pathways and send projections to corresponding cortical areas. Axons from TC cells also collateralize upon cells of the thalamic reticular nucleus (TRN) *en route* to the cortex (Lam and Sherman, 2011; Scheibel and Scheibel, 1966). The TRN is a shell of GABAergic neurons that surround the thalamus in its dorsal aspect (Pinault, 2004). Neurons of the TRN also receive excitation from corticothalamic cells of layer V and VI (Bourassa and Deschênes, 1995; Carroll et al., 2022; Whilden et al., 2021), and a wealth of modulatory input (Pazo et al., 2013; Herrera et al., 2016; Beierlein, 2014; McCormick & Wang, 1991; Aizenberg et al., 2019). In light of its potential to selectively inhibit sensory relay, TRN has been proposed to focus an attentional “searchlight” on behaviorally important information before it reaches the cortical input layer (Crick, 1984; McAlonan et al., 2008). A variety of behavioral and recording experiments have demonstrated the influence of TRN towards attentional tasks (Weese, 2022; Wells et al., 2016; Wimmer et al., 2015; McAlonan et al., 2000; McAlonan et al., 2006; Nakajima et al., 2019). Anatomical data also support potential intra-TRN and feedback connections that may implement the searchlight (Pinault and Deschênes, 1998a, 1998b; Pinault et al., 1995; Shosaku et al., 1989; Scheibel and Scheibel, 1966). Yet the specific influences of TRN circuits on thalamocortical relay have not been well described.

Communication between TRN cells is proposed as a key feature of the proposed spotlight function in thalamocortical circuits (Pinault and Deschênes, 1998a). Whether the GABAergic neurons of the TRN chemically synapse onto each other remains controversial. Inhibitory responses have been recorded from glutamate uncaging experiments in juvenile mice (Deleuze and Huguenard, 2006; Lam et al. 2006), but optogenetic stimulation of ChR2-expressingTRN cells revealed that direct inhibitory synapses appear to be limited to earlier development (Hou et al. 2016). Only a single direct demonstration of a mixed inhibitory and electrical connection between paired TRN cells has been reported (Landisman and Coulon, 2024). Connectivity within TRN is instead mainly thought to rely on its dense population of electrical synapses (Landisman et al., 2002). Coupling between TRN cells appears to be extensive and may couple up to 20 cells with the average coupled network size being 9 cells, as revealed by dye-coupling experiments (Lee et al., 2014). Electrical synapses can play many diverse roles within networks; the type most often described and well-understood is enhancement of synchrony amongst coupled neurons (Christie et al., 2005; Lewis and Rinzel, 2003; Leznik and Llinás, 2005; Mancilla et al., 2007; Pfeuty et al., 2005). TRN activities are more synchronous when electrical synapses are in a stronger state (Long, 2004). Beyond their synchronizing influence, recent large scale simulations of TRN, which included electrically coupled networks have often focused on aspects of rhythmicity or sleep spindle activity (Iavarone et al., 2023, Li et al., 2024). Electrical synapses of the TRN may exert other complex effects towards processing transient inputs; models have shown that the strength of coupling can merge closely timed inputs or act to further distinguish inputs with greater difference in timing or amplitude (Pham and Haas, 2018).

How electrical synapses impact the passage of transient inputs within TRN or across the brain’s networks is not well understood. Previous demonstrations of transient signal processing by electrical synapses have focused on very local interactions. Coincident inputs received by two coupled cells can be enhanced through the presence of electrical synapses (Galarreta and Hestrin, 2001 Veruki and Hartveit, 2002). Similarly, temporally separated inputs to two neurons become weaker due to the current sink produced by a coupled quiescent cell (Trenholm et al. 2013; Alcami, 2018). Transmission of inhibitory afterhyperpolarizations by electrical synapses in a rhythmic context acts to desynchronize cell activity (Vervaeke et al., 2010; Hurkey et al., 2023). Coupling between cerebellar basket cells increases spiking probability in those cells through spikelet transmission, evoked within a short time window (Alcami, 2018; Hoehne et al., 2020). Similarly, electrical synapses enhance synchrony and also temporal precision in the toadfish vocal circuit (Chagnaud et al., 2021). Chemical EPSPs transmitted along a dendrite can be shunted by the presence of nearby electrical synapses (Lang et al., 1996; Llinás et al., 1974; Mendoza and Haas, 2022). These effects may also act to enhance signal-to-noise ratio in individual neurons through the combined effects described above (Ammer et al., 2022). In canonical feed-forward circuits electrical synapses between interneurons broaden the integration window of target cells and can enhance transmission efficiency in larger networks (Pham and Haas, 2019).

We sought to understand how electrical synapses embedded within a network of TC and TRN neurons regulate the processing of ongoing sensory inputs during relay to cortex. Working from our previous model that used a single electrical synapse between two TRN neurons to reveal modulation of the temporal dynamics of TC cell responses (Pham and Haas, 2018) before they are relayed to cortex, we constructed a coupled network of a ring of 9 TRN and 9 TC cells receiving a sequence of synaptic inputs, such that each subsequent TC response is modulated by the prior responses of its neighbors. We added a single cortical integrator neuron to this model in order to examine the ultimate impacts of TRN electrical synapses upon thalamocortical relay. This architecture allowed us to examine how synapses within TRN modulated responses of the TC cells, and the information that is relayed to cortex. Our simulations showed us that spiking properties and importantly, correlation and separation of spikes in the thalamocortical relay were altered as a function of coupling strength, highlighting the complex and pervasive roles of electrical synapses in ongoing activity as it passes through networks. These results also provide a basis for understanding how plasticity of electrical synapses affect the networks in which they are embedded.

## Results

Our general goal was to investigate the reach of electrical synapses; that is, within a network that includes coupled neurons, the radius of their functional influence. First, we measured coupling coefficients between TRN cells using the commonly used experimental paradigm of delivering a hyperpolarizing pulse to one cell and measuring voltage deflections through the rest of the network (Figure 1A, top). For the strongest synapses, a pulse that revealed a coupling coefficient of 0.2 in the neighboring cell diminished to 0.05 at the next cell and diminished to negligible thereafter (Figure 1B). These coupling coefficients suggest minimal influence on the voltage signals extends beyond at most 3 synapses from the originating neuron even for high electrical synapse strengths. We used the same network to compare responses to identical inputs to each of the neurons (Figure 1A, bottom), with a delay of 20 ms between inputs. In this scenario, we noted that responses of each sequential cell in the chain included more and/or faster spikes, resulting in a cumulative rather than diminishing effect on spiking in each neighboring neuron (Figure 1C). This exercise demonstrates that coupling coefficients may not capture or predict the impact of electrical synapses within active neuronal networks.

**Figure 1.**
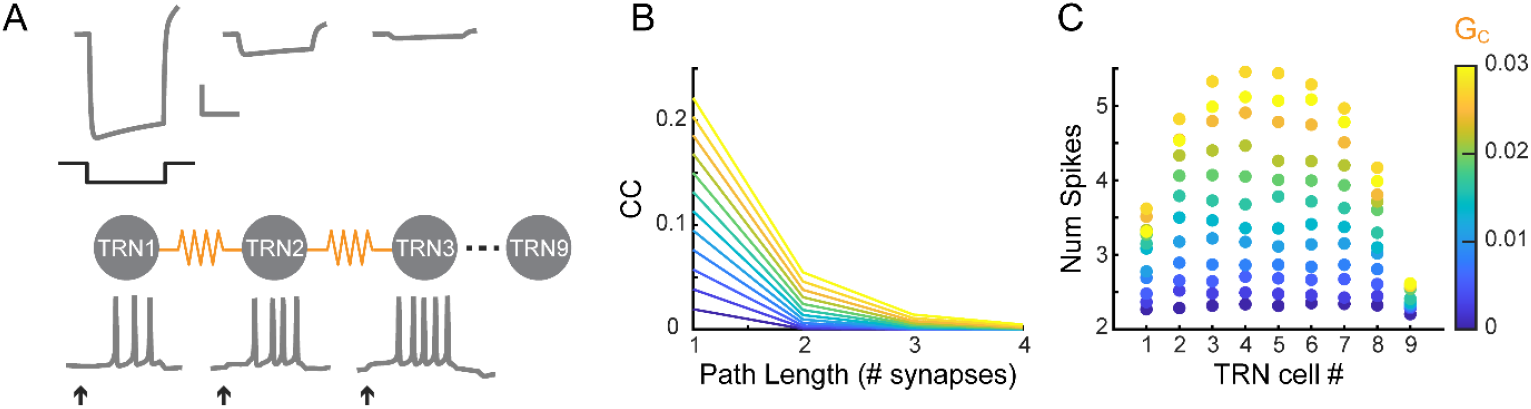
Effects of electrical synapses measured over network distance. A). Simple network of identical coupled TRN neurons. Top traces: voltage responses to a hyperpolarizing current applied to TRN1. Bottom traces: spiking responses to identical excitatory inputs (arrows) delivered to each cell. B). Coupling coefficients (cc) as a function of path length across the TRN network in A, as measured from TRN1. C). Average number of spikes in the response of each TRN cell in the network resulting from inputs shown in A.

We then set out to address how electrical synapses embedded within a network of TRN neurons regulate the processing of ongoing sensory inputs relayed by a layer of thalamocortical (TC) relay neurons. For this network, we provided a sequence of synaptic inputs to the TC cells of a 9-cell ring (Figure 2A). All cells were provided with spontaneous input such that they fired spontaneously at 6 - 8 Hz (Figure 2B); this choice rendered them more sensitive to smaller additional inputs we provided, compared to neurons at rest that would require larger inputs. Synaptic inputs were single falling-exponentials with a 30-ms decay time constant. We chose a conductance for the synaptic inputs that, in the absence of other connections within the circuit, drove a 4-spike burst in the TC neurons at a latency of ∼15 ms (Figure 2B). We used excitatory chemical synapses between TC and TRN neurons strong enough to drive a 3-spike burst in the TRN neurons at a latency of ∼30 ms from the input. Inputs were delivered at a fixed time difference (Δt_in_), between 10 and 50 ms; we chose this set of inputs to represent a spectrum of thalamic inputs ranging from nearly identical to less similar (e.g. whiskers moving across a surface; similarly timed or frequencies of sound). Because coupling introduces a current sink to a neuron that reduces input resistance (Amsalem et al. 2016) we adjusted leak conductances current in order to maintain TRN input resistances at a constant value as we varied coupling across simulations. These choices and input sequence gave us a basis for evaluating the impact of electrical synapses between TRN neurons, whose effects were exerted through the inhibitory synapses from each TRN cell to its paired TC cell.

**Figure 2.**
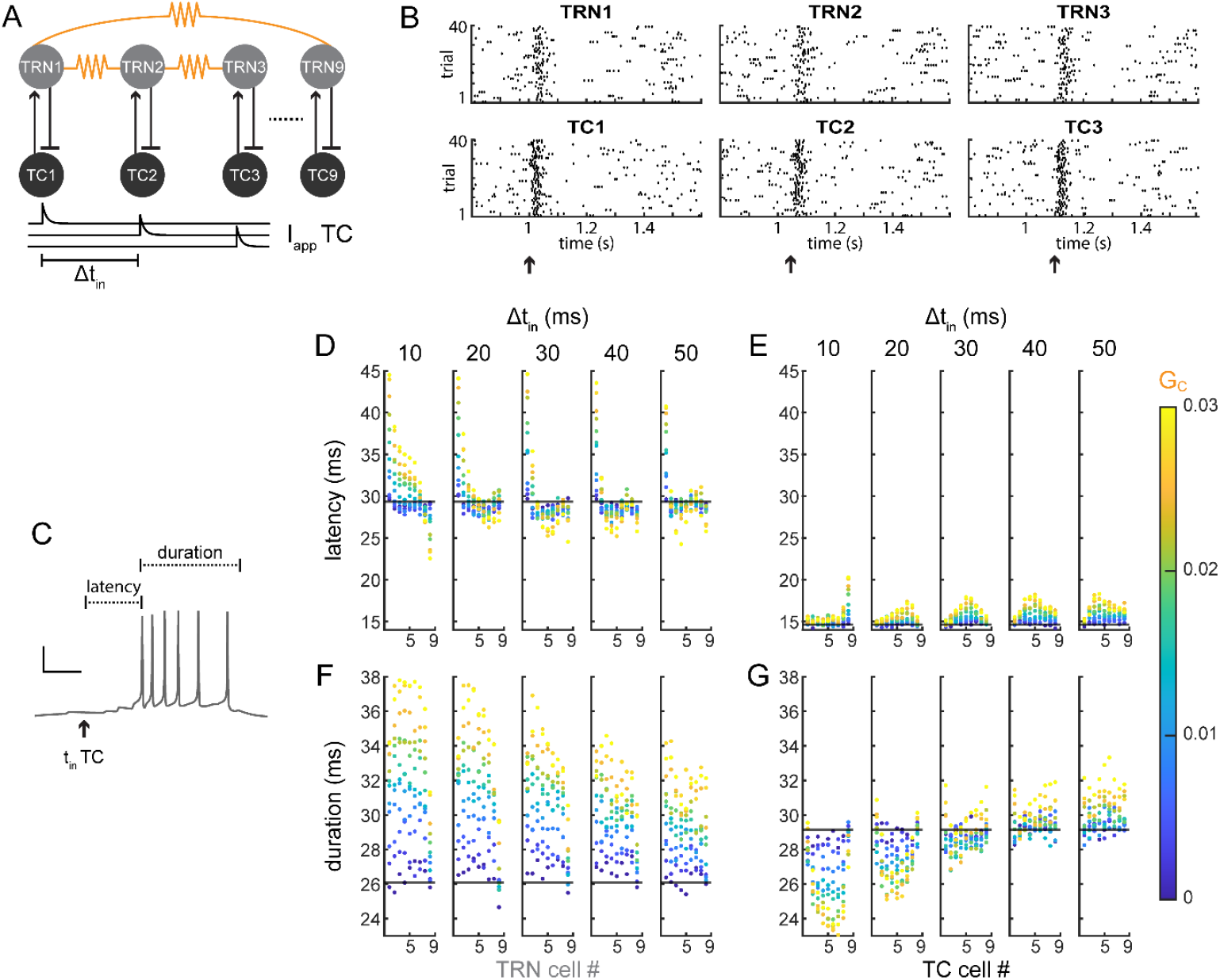
Coupling within the TRN layer modulates the latency and duration of TRN and TC spike trains. A). Schematic of the ring network and inputs. B). Example rasters of TRN and TC spike trains. C). Representative trace showing measurements of spiking properties resulting from an input delivered to one TC cell. Scale bars: 25 mv, 20 ms. D). Average spiking latency of TRN cells in the network with increasing G_c_. Black lines represent latency in an uncoupled network. Left to right panels: increasing Δt_in_. E). Average spiking latency of TC cells with increasing G_c_ within the TRN cell network. Black lines represent values in an uncoupled network. Left to right panels: increasing Δt_in_. F). Average spiking duration of TRN cells in the network with increasing G_c_ value. Black lines represent duration in an uncoupled network. Left to right panels: increasing Δt_in_.G). Average spiking duration of TC cells with increasing G_c_ within the TRN cell network. Black lines represent duration in an uncoupled control network. Left to right panels: increasing Δt_in_.

To understand the impacts of electrical synapses within the TRN layer on TC spiking, we first measured the latency and duration of spiking responses in the TRN cells as a function of electrical synapse strength and input timing (Figure 2C). In the TRN neurons, increases in electrical synapse strength increased the latency of TRN spiking (Figure 2D), consistent with increases in leak conductance. This effect was proportional to electrical synapse strength and was largest for TRN1, which experienced an increase from 30 to 45 ms, and diminished by TRN9. Latency changes were largest for more closely-timed inputs. The duration of TRN spike trains was also modulated by electrical synapses, with consistently longer duration proportional to coupling strength and slightly diminished with increased Δt_in_ (Figure 2F). Overall, effects of electrical synapses within TRN cells were strongest for the several neurons closest to each synapse, as is intuitive.

Next, we turned to the TC layer. Because inputs arrived to TC cells first, latency of spiking was only minimally longer, less than 5 ms, in TC cells with greater TRN coupling strength (Figure 2E). Those increases were a result of increased spontaneous activity within the TRN layer mediated by increased GJ strength; that activity then resulted in increased inhibition delivered to TC cells. Duration of TC spiking (Figure 2G) was more strongly controlled by the inhibition delivered from TRN cells, which was in turn modulated by electrical synapses between TRN neurons. As a result, TC spike trains were truncated by up to 5 ms when both coupling was high between TRN cells and Δt_in_ was small (Figure 2G, left). However, coupling also produced an increase in spiking duration by several milliseconds in TC cells when Δt_in_ exceeded the duration of TC spiking (Figure 2G, right). Overall, we note that the effects of TRN synapses accumulated in the TC layer with time difference and distance. This result highlights that coupling within the TRN network produced differential effects on TRN and TC layers that were dependent on input time difference in the TC layer and on coupling strength.

Electrical synapses in the TRN layer also modulated the correlations of TC and TRN neurons to the input. We measured correlation between the input times and spike trains in the coupled network, each convolved by a falling exponential with time constant 5 ms. TRN cells were least correlated to inputs for closely-timed inputs and strong electrical synapses (Figure 3B, left), diminishing by 50%, as a result of the GJ-driven changes in latency and duration of their spike trains described above. A cumulative effect of electrical synapses can be seen across TRN cells, producing a gradual increase in correlation from the first to last TRN to receive the input. Correlation of TC cells to their inputs was also modulated, although much less strongly, by GJs between the TRN neurons, and again in a cumulative manner across TC cells (Figure 3C). These effects were driven mainly by changes in duration of TC spiking, which differed most greatly at the extremes of Δt_in_ and produced the lowest correlations at those values as well.

**Figure 3.**
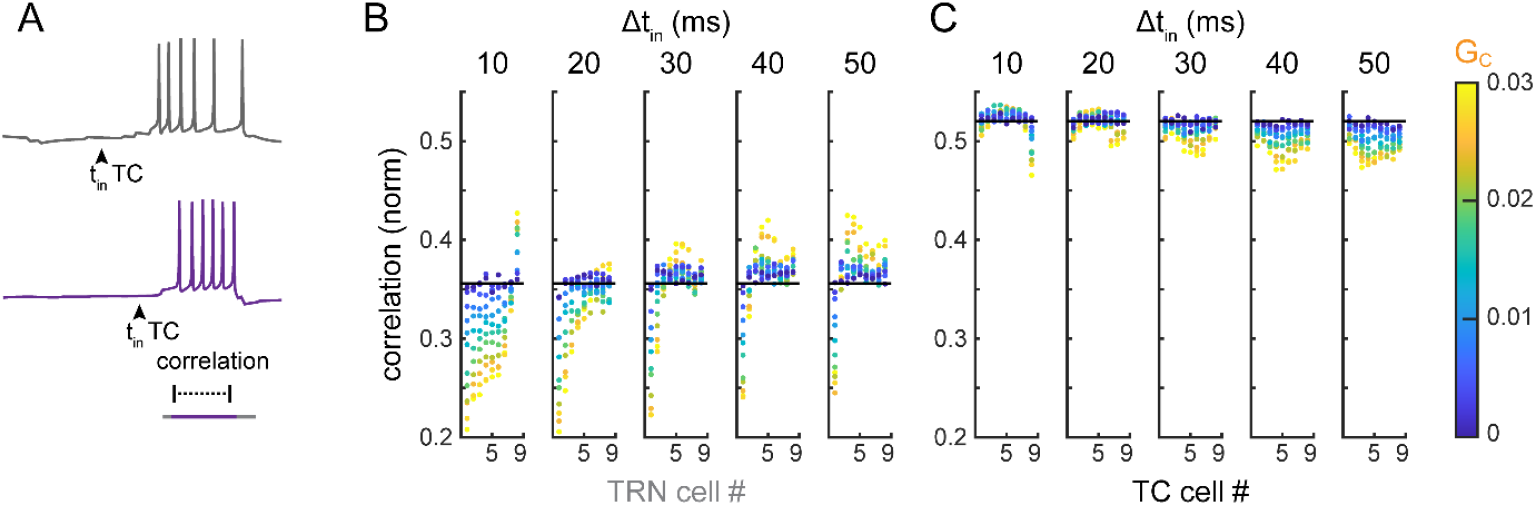
Electrical coupling decorrelates TRN and TC trains from sensory inputs. A). Representative trace showing measurement of correlation between spike trains from an uncoupled network (gray) and GJ-modulated spike trains (purple). B). Average correlation between input and TRN cells for increasing G_c_ value. Black line represents correlation for an uncoupled network. Left to right panels: increasing Δt_in_. D). Average correlation between input and TC cells for increasing G_c_ within the TRN layer. Black line represents separation correlation in an uncoupled network. Left to right panels: increasing Δt_in_.

As a first step at understanding how cortical reception of thalamocortical relay is modulated by TRN electrical synapses, we measured the separation and overlap between TC spike trains (Figure 4A) (Pham and Haas, 2018). Greater separation between TC spike trains potentially underlies discrimination by a cortical neuron that is integrating TC inputs. As a result of inhibition from the TRN spike trains, coupling strength altered the separation values of each pair of TC spike trains. When Δt_in_ was small, TC spikes had significant overlap (separation < 0) in the uncoupled networks; increasing electrical synapse strength produced increased separation and reduced overlap (Figure 4B, left). At longer Δt_in,_, whereTC spikes were already positively separated, the overall separation in TC spike trains was reduced by greater electrical coupling (Figure 4B, right). In both cases, the effects increased across pair number as more inputs were delivered to the TC layer, again demonstrating a cumulative effect due to network coupling (Figure 4C). The magnitude of separation changes was considerable at the timescale of spike timing, as the effect of coupling produced up to 10 ms difference from control spike separations (Figure 4D). In summary, the decorrelation of TC cell responses, and thus changes in how separated responses were from control, are produced by both latency and duration changes dependent on the coupling within the network. The consequences of these temporal changes predict substantial changes in how the cortex might integrate and process the arrival of these thalamocortical inputs.

**Figure 4.**
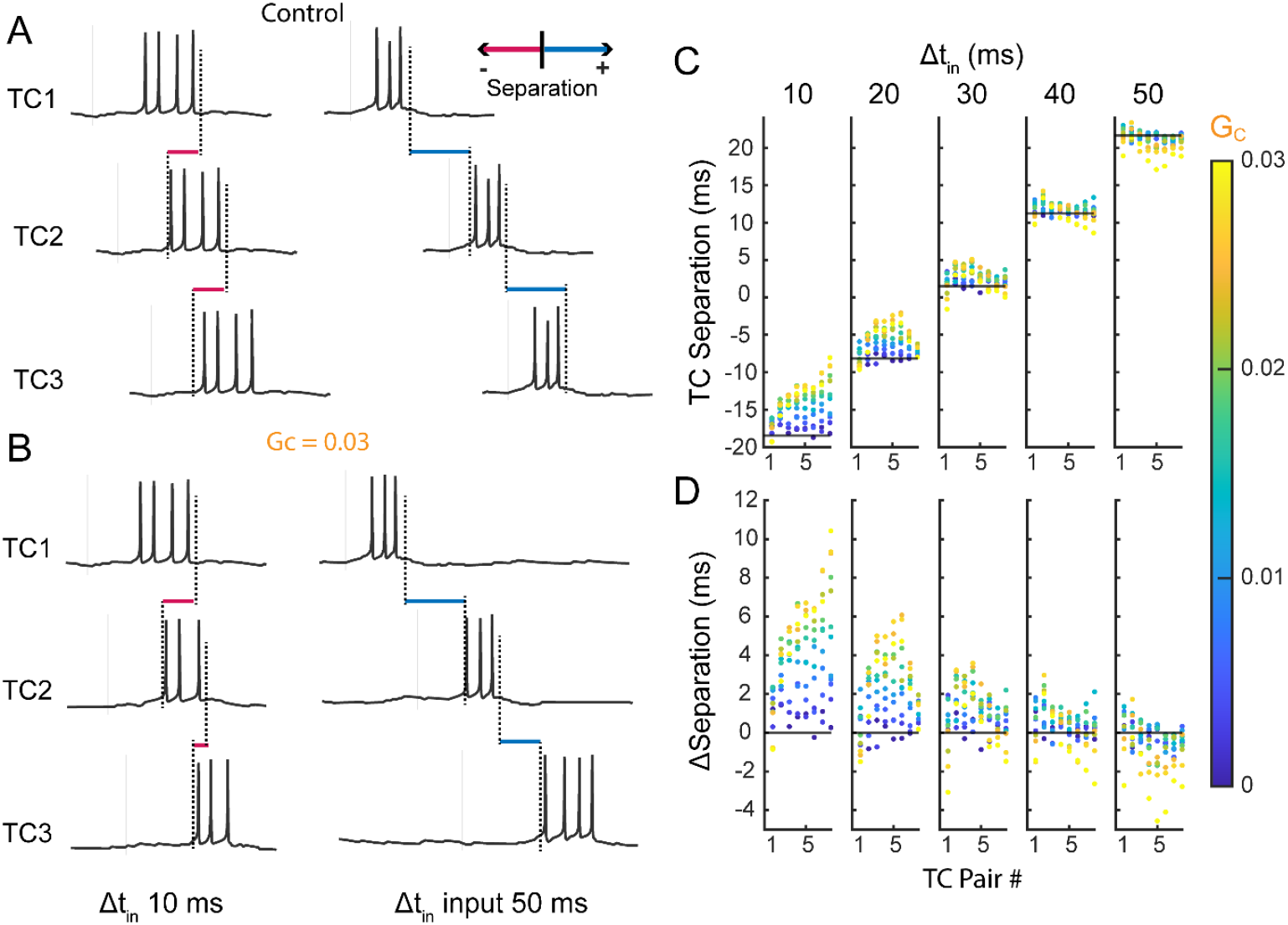
Coupling within TRN layer modulates separation of TC spike trains. A). Representative traces from the first three TC cells for a control trial (no coupling between TRN cells). Left traces show separation with a small Δt_in_, Right traces show separation with a large Δt_in_, with separation notated. B). As in A, for a trial with high coupling strength between TRN cells C). Average pairwise spike separations of TC cells in the network with increasing values of uniform G_c_ between TRN cells. Black lines represent separation in an uncoupled network. Left to right: increasing Δt_in_. D). Average pairwise changes in spike separation of TC cells, plotted for increasing G_c_ within the TRN cell network, relative to the control response of TC to input alone (black lines). Panels show results for increasing Δt_in_.

Finally, we added a single simple integrator neuron to sum and read out the output of thalamocortical relay from our ring network (Figure 5A). Because spontaneous TC spikes could confound or mask the effects of electrical synapses within the TRN layer on cortical integration, we ran these simulations in the absence of noise. Instead, to produce variation across trials, we delivered inputs with 5% variation in the ISI. We chose an excitatory synapse strength for TC inputs to the cortical neuron that produced a single spike in the cortical neuron for each TC input (Figure 5B, bottom). Cortical spike times were nearly regular (CV of spike trains = 20-30%) for no or low TRN coupling. Surprisingly, we observed that increases in coupling within the TRN layer produced more spikes, irregular spikes within trains, and longer spike trains in the cortex cell across all Δt_in_ values (Figure 5B). Interspike interval (ISI) distributions for cortical spikes were tightly restricted around the Δt_in_ value in the uncoupled and low-coupling ranges (Figure 5C, bottom). With increased coupling strength within the TRN, ISI distributions in the cortical neuron broadened substantially (Figure 5C, top). To better visualize this effect, we plotted the gain of the cortical ISI distributions in coupled cases relative to the ISI distributions for uncoupled TRN (Figure 5D). These plots revealed a reduction in ISIs around the value of Δt_in_ (dotted line) and a gain in shorter ISIs as coupling strength increased, indicative of the shift from regular spiking towards more irregular firing. Consistent with the increase in shorter ISIs, the average spiking output of the cortex cell increased for higher coupling within TRN (Figure 5E). We quantified the effect on irregular firing by measuring the coefficient of variation (CV) of the ISI distribution. Spike trains in cortex were increased in CV to 60% at higher coupling values and across Δt_in_ (Figure 5F). Lastly, we asked if these spiking changes modulated the correlation of cortex spikes to the TC input train. Correlation in the cortex spikes to the input train was high (70-90%) for uncoupled and weakly coupled TRN and decreased (to 40-50%) as a result of increased coupling (Figure 5G). We conclude that electrical synapses within the TRN have diverse impacts on TRN spiking, which accumulated and propagated to modulate TC cell responses, and ultimately, cortical responses to thalamocortical inputs.

**Figure 5.**
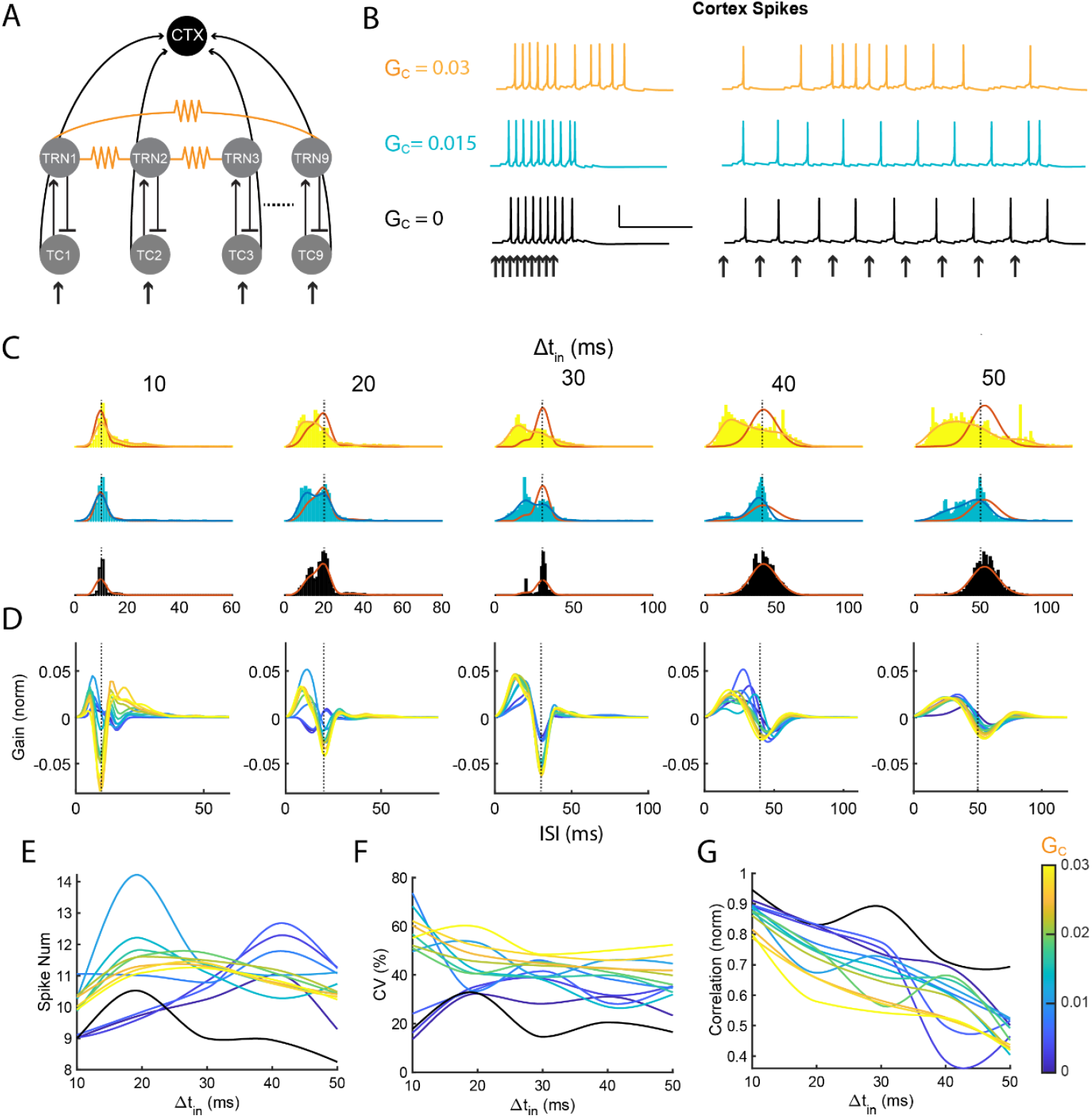
Impact of coupling within TRN on TC-relayed cortical spiking. A). We added a cortical neuron to integrate TC relay from the ring network. B). Cortical spike trains for control (no coupling between TRN cells), moderate (blue), and high (yellow) coupling between TRN cells within the network, for. Δt_in_ = 10 ms (left) and Δt_in_ = 50 ms (right). C) ISI distributions for the cortex cell for control (no coupling between TRN cells), moderate (blue), and high (yellow) coupling between TRN cells of the network. Mean ISI distributions for uncoupled TRN are overlayed in red. Left to right: increasing Δt_in_. D). Gain of the ISI distributions, versus time and across coupling values. E). Average number of spikes in the cortex cell, versus Δt_in_ and across coupling values. F). Average coefficient of variation of the ISI distributions in C, versus Δt_in_ and across coupling values. G). Average correlation between cortex spike times and input sequence times, versus Δt_in_ and across coupling values.

## Discussion

While the role of electrical synapses in synchronizing rhythmic activity is well established, their contributions to the neuronal processing of transient inputs across networks remains less well characterized. Our computational investigation demonstrates that embedded electrical synapses have an unexpectedly decorrelating and potentially strong impact on thalamocortical relay that arises as the modulation of electrical synapses between TRN neurons propagates and accumulates through the network. Electrical synapses between TRN cells modulate latency and especially duration of TRN spike trains; these changes are passed into the TC cells through inhibitory synapses and impact the duration and correlation of TC spiking. The effects of each electrical synapse within the chain accumulate with each cell, and result in dispersed and decorrelated responses in the cortical cell. These effects illustrate that TRN coupling influences activity through a large extended network, with effects on spiking output that would not be predicted from the passive measurements of each electrical synapse alone, as coupling between pairs alone would not predict the effects we see in a network context and drops to indistinguishable levels when measured by the traditional hyperpolarizing current steps across multiple synapses.

TRN cells couple small clusters of nearby cells with direct electrical synapses; our model here considers a specific topology of connections in order to address the extent of electrical synapse influence over spike times. Effects of gap junctions on transient inputs, which note excitatory effects for closely timed inputs, would also propagate across several cells in a network context and have considerable impacts on information processing (Pham & Haas, 2018; Trenholm et al., 2013; Hoehne et al., 2020; Vervaeke et al., 2010; Galarreta & Hestrin, 2001). Our simulations provide basic investigations into larger electrical synaptic embedded networks at TRN, which can form a basis of comparison for future investigations. In particular, whether these results can be reproduced in a more naturalistic topology in which connections are randomly dispersed among nearby cells, remains to be seen. Plasticity of electrical synapses, despite occurring over timescales much larger than our transient input scheme, would modify activity of ongoing sensory relay in the long term. Numerous mechanisms of plasticity have been described in coupled systems (Haas et al., 2011; Lasater and Dowling, 1985; Mathy et al., 2014; Pereda and Faber, 1996); Landisman, 2005; Wang et al., 2015). Though we did not include plasticity dynamics in our model, the effects of modifying synapse strengths within the network can be extrapolated from the synapse strengths we sampled in our network model to provide specific predictions.

Many refinements are possible for our network model that could impact its predictions. Our thalamic cell models were detailed in order to include the important bursting behaviors in those cell types (Destexhe et al., 1998; McCormick & Huguenard, 1992; Destexhe et al., 1996), but we opted for a simplistic cortical output neuron so that changes we observed in responses were due to the electrical synapse-mediated changes in the relay alone. We expect that intrinsic properties of cortical cells could be another important source of modulation in cortical response that would interact with the modulation by TRN synapses. Thus, further study could include more detailed neuron models for layer 4 and include specifics such as spike adaptation due to M-currents, or dendritic morphology and dynamics. Further, deep-layer cortical neurons provide an important source of input to TRN (Hádinger et al., 2023; Carroll et al., 2022; Gentet & Ulrich, 2004; Whilden et al., 2021; Sokhadze et al., 2025), and the impacts of that connection combined with electrical synapses within TRN should also be expected to modulate thalamocortical transmission. Also absent from our models are lateral inhibitory connections between nearby TRN cells (Deleuze and Huguenard, 2006; Lam et al., 2006, Landisman and Coulon, 2024). Corticothalamic feedback onto both TC and TRN neurons, although delayed, could be another avenue of investigation for future experiments. Another key thalamic synapse proposed as a mechanism of the searchlight is open-loop divergent inhibition from TRN to TC cells (Crabtree et al., 1998; Crabtree and Isaac, 2002).

TRN has recently been shown to comprise distinct cell types which differentially target thalamic relays of first and higher-order information (Clemente-Perez et al., 2017; Martinez-Garcia et al., 2020). Our results here only consider one homogenous source of TC relay, and do not include the influence of these cell types and the potential sensory modulation that higher order sources may contribute (Bu et al. 2025). Electrical synapses have been shown to couple TRN cells across cell type (Vaughn et al. 2024), additionally SOM+ TRN may provide lateral inhibition to the first order PV+ TRN cell type. In all, our model can provide important predictions for the diverse connections both within and between thalamic sources and their impact on precise sensory relay to the cortex.

## Methods

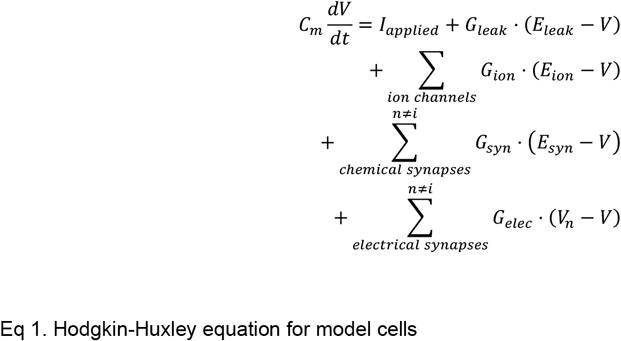

Eq 1. Hodgkin-Huxley equation for model cells

Models for all cells in were Hodgkin-Huxley point models (Eq. 1) were written in the Julia programing language and solved using the DifferentialEquations.jl ODE solving suite (Rackauckas and Nie 2017). Parameters for TC and TRN cells were built upon those used previously (Traub et al. 2005, Destexhe et al. 1994a, Pham and Haas, 2018), and included the following ionic currents and maximal conductances: fast transient Na+ (NaT) 60.5 mS/cm^2^, K+ delayed rectifier (Kd) 60 mS/cm^2^, K+ transient A (Kt) 5 mS/cm^2^, slowly inactivating K+ (K2) 0.5 mS/cm^2^, slow anomalous rectifier (AR) 0.025 mS/cm^2^, and low threshold transient Ca2+ (CaT) 0.75 mS/cm^2^.Reversal potentials were 50 mV for sodium, −100 mV for potassium, 125 mV for calcium, −40 mV for AR and −75 mV for leak. Capacitance was 1 µF/cm^2^ with leak of 0.1 mS/cm^2^. Cortical output cell was a regular spiking neuron with only regular sodium and potassium currents and a leak of 0.1 mS/cm^2^ (Pospischil et al., 2008). Chemical synapses were modelled as double exponential decay with rise and fall time kinetics of 5 ms and 35 ms respectively, with reversal potentials of 0 mV for excitatory and −100 mV for inhibitory synapses. Electrical synapses were modelled as static ohmic conductance applied to the voltage difference between the coupled TRN cells.

Scaling up to a network, the excitability changes as coupling is increased in a network (Amsalem et al. 2016). Thus, for each coupling strength, TRN cell leak conductance was systematically scaled down to match the input resistance of the uncoupled TRN model. Sensory inputs were simulated as exponentially decaying current pulses (peak amplitude 0.75 µA/cm^2^, with decay time constant 30 ms) delivered simultaneously to each of the TC cells. Chemical synaptic connections between TRN cells were set at 1 µA/cm^2^ for excitatory synapses from TC to TRN cells, and 1 µA/cm^2^ for all inhibitory synapses originating from TRN. For each stimulus condition, we repeated 1500 trials under Poisson random noise at a rate of 80 Hz for excitatory and 20 Hz for inhibitory events. Coupling also increased the baseline spontaneous firing rate of the TRN neurons, therefore we scaled the amplitude of noise to maintain all cells at a spontaneous firing rate of 6-8 Hz across all simulations. Subthreshold current steps were applied to all cells to bring cells close to threshold, TC and TRN cells were depolarized to −70 mv, cortical cells were given 0.3 µA/cm^2^. Spike train data was extracted and saved for further analysis and visualization using MATLAB.

### Spike train analysis

Separation of TC cells spikes were calculated as the time difference between the last spike of the first TC cell pair to receive the input, and the first spike of the following cell response. Correlation was calculated from overlap of input time and spiking response time series, convoluted by a falling exponential to produce similar time series to the delivered input. Latency was simply the difference between input onset and the time of the first spike in the response. Duration was the difference between the first and last spike of the response, responses consisting of only one spike were excluded from analysis. Trials consisted of 1500 replications under Poisson random noise and used to calculate mean and standard errors of response variables. If no spike occurred within 100 ms following an input the trial for that cell was excluded from calculated means and standard errors for those response characteristics.

## Code Availability

All code used is available through Github. The code/software can be found at: https://github.com/jhaaslab/RingModel

